# Magnetic resonance imaging of the regenerating neonatal mouse heart

**DOI:** 10.1101/329474

**Authors:** Mala Gunadasa-Rohling, Megan Masters, Mahon L. Maguire, Sean C. Smart, Jürgen E. Schneider, Paul R. Riley

**Affiliations:** British Heart Foundation Oxbridge Centre of Regenerative Medicine, Department of Physiology, Anatomy and Genetics, University of Oxford, Oxford, OX1 3PT, UK.; British Heart Foundation Experimental Magnetic Resonance Unit, Wellcome Trust Centre for Human Genetics, Roosevelt Drive, Oxford OX3 7BN, UK; RDM Cardiovascular Medicine, John Radcliffe Hospital, Headley Way, Headington, Oxford OX3 9DU, UK; Cancer Research UK and Medical Research Council Oxford Institute for Radiation Oncology, Department of Oncology, University of Oxford, Oxford, United Kingdom; Division of Biomedical Imaging, Leeds Institute of Cardiovascular and Metabolic Medicine, LIGHT Laboratories, Clarendon Way, University of Leeds, Leeds, LS2 9JT, UK

**Author notes:** Joint senior author; correspondence to and.

## Abstract

We present longitudinal magnetic resonance imaging (MRI) of neonatal mouse hearts during the first three weeks following coronary artery ligation to mimic heart attack. We confirm heart regeneration in individual animals injured on post-natal day 1 (P1) while those injured on P7 show the adult response of fibrosis, scarring and impaired heart performance. We document heart growth and development of the principal functional cardiac parameters, and also remodeling during tissue regeneration as compared to fibrosis when imaging repeatedly up to 21 days after myocardial infarction (MI). We reveal compensatory changes in cardiac function with the restoration of tissue and resolution of injury for the P1 cohort and sustained injury responses for the P7 cohort. This study resolves the controversy surrounding neonatal mouse heart regeneration and establishes a functional platform for live capture of the regenerative process and for the future testing of genetic or therapeutic interventions.

---

Human patients who suffer a myocardial infarction (MI or “heart attack”) are unable to replace the lost or damaged heart muscle (up to 25% of the myocardium) and instead a non-contractile fibrotic scar is laid down at the site of injury to prevent cardiac rupture. The result is compensatory function and pathological remodeling of the surviving myocardium (irreversible chamber dilation and wall thinning) which ultimately leads to congestive heart failure, the major contributor to morbidity and mortality worldwide^1^. Strategies in regenerative medicine to replace lost myocardium include stimulating resident heart muscle cell (cardiomyocyte) division, direct reprogramming of fibroblasts into cardiomyocytes, and transplantation of cells that may either transdifferentiate into new heart muscle or, through paracrine effects, promote repair and regeneration (reviewed in^2^). In addition, studies of animal models which can intrinsically regenerate injured myocardium, such as the adult zebrafish^3^, have proven invaluable in identifying cellular and molecular mechanisms underlying this regenerative capability. In 2011, the first evidence of mammalian heart regeneration was reported by Hesham Sadek, Eric Olson and colleagues^4^, Following surgical resection of ∼15% of the left ventricle apex of a one-day old (P1) neonatal mouse, the heart fully regenerated by 21-days post-injury, whereas if the procedure was repeated one week later on a post-natal day 7 (P7) mouse heart, fibrosis and scarring ensued, recapitulating the adult wound healing response. The mechanism of regeneration observed was analogous to that described in the adult zebrafish heart^5, 6^, in the P1 mouse there was evidence of significant cardiomyocyte turnover and proliferation to restore damaged heart muscle, which was lost by P7. Since the original study several others have described neonatal myocardial regeneration after resection and following alternative insults, such as cryo-injury^7^, and coronary artery ligation to invoke MI^8^. Others have documented roles for specific cell types such as tissue resident macrophages^9^ or implicated important regulatory pathways such as Hippo signaling^10^ in neonatal myocardial regeneration. Despite the apparent utility of the various models of neonatal heart injury, and the resulting tissue response, there is some controversy surrounding the extent of regeneration during the first weeks of life. It has been reported that, following apical resection, regeneration did not occur and was replaced by a fibrotic response^11^ which persisted long-term (180 days) with extensive cardiac remodelling^12^. The differences between the results of Andersen et al. and those of the original report^4^ were not due to genetic background, instead differences in the method of injury (i.e. removing up to 40% of the ventricle as opposed to ∼15%) were suggested to be incompatible with regeneration^13, 14^.

The major issue to-date with all of these studies is that they are based on sacrificing individual animals at specific time-points post-injury followed by histological assessment of the heart. This provides no insight into the extent of the initial injury following surgery, whether the heart was indeed injured from the outset or the regenerative process over time. Moreover, there remains an important question as to how the regenerating mammalian heart copes in terms of functional output and remodeling during the processes of scar resolution and tissue restoration, as is directly relevant to human ischemic heart disease patients subjected to regenerative therapies. To address both of these key issues, we have developed a non-invasive Magnetic Resonance Imaging (MRI) approach that enabled tracking of individual new-born mice from the first day after birth over time, and also after myocardial injury. While MRI is well established for quantifying and monitoring cardiac function in adult mice^15, 16^, its longitudinal application in neonatal mice requires significant refinement, and has not yet been reported at the early P1 time point, or following neonatal heart injury.

## Results

### Growth and development of the neonatal mouse heart and early cardiac function parameters

The non-invasive nature of MRI allowed us to track the growth and developing cardiac function in the same neonate over time starting from P1, with five scans performed over a period of three weeks (day 0 – baseline, days 4, 7, 14 and 21 after injury/sham-operated controls), corresponding pup ages of P1, P5, P8, P15 and P22. A similar analysis was carried out for our P7 cohort, providing data for P7, P11, P14, P21 and P28. The mouse pup was positioned in a modified MRI cradle for imaging (Supplementary Figure 1) and scanned for 35-45 minutes each time. The values of left ventricular mass to body mass ratio (LVM/BM), heart rate (HR) and ejection fraction (EF) for the sham-operated animals (which serve as our control group) are depicted in Figure 1, along with a body mass (BM) growth chart. BM increased linearly until day 21, after which the rate of increase was higher due to the pups being weaned. With recurrent sampling, we observed an increase in the growth rate of the P7 group vs the P1 group (Fig. 1a). It has been suggested that repeated administration of anesthesia may have a negative effect on body growth^16^, consistent with this the P1 group that received anesthesia and scanning earlier weighed less than the P7 group at three comparable time points (P7-8, P14-15, and P21-22). The mean weights for the P1 group were nearly 2.5g lighter than the P7 at the midpoint, though they appeared to recover with age.

**Figure 1.**
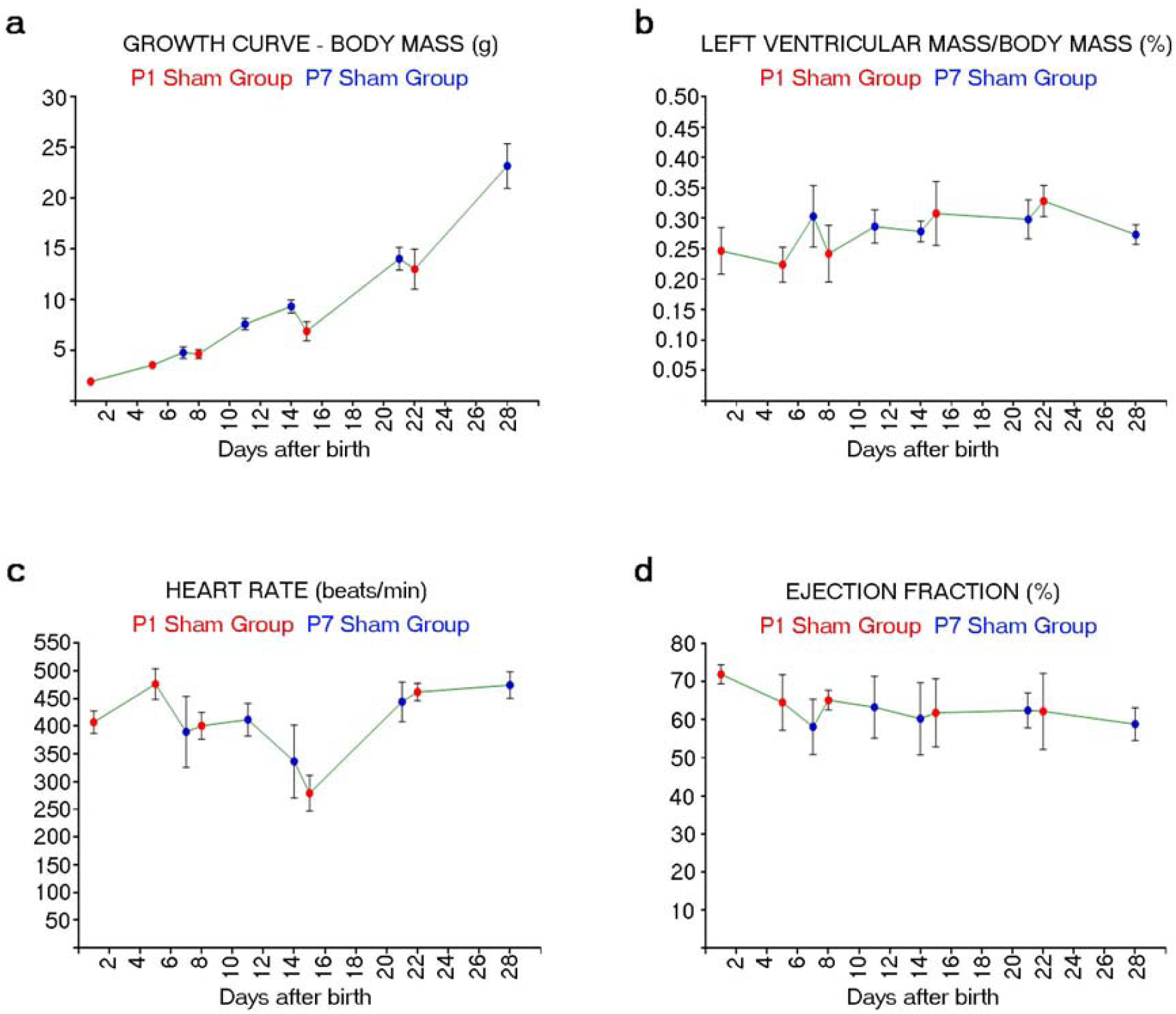
Growth and development of the neonatal mouse heart. Development of (a) body mass (BM), (b) left ventricular mass to body mass ratio (LVM/BM), (c) heart rate (HR) and (d) ejection fraction (EF) in sham control animals over the study duration of 28 days; n=6 for all groups.

Left-ventricular mass index (i.e. LVM normalized to body mass), increased over time within the P1 group (Fig. 1b), which is contrary to a previous study^15^ that reported a decrease in LVM/BM with age. The differences between our findings and those previously reported^16^ relate to the stages examined, in that we imaged early time points throughout the first three weeks of life when heart growth rate is maximal, whereas the previous study assessed older-aged animals of up to four months where the relative increase in LVM is reduced. The P7 group did not show this early increase and in fact revealed a steady reduction over time in LVM.

HR increased from P1 to P5 (Fig. 1c), likely caused by an increasing anesthesia tolerance of the maturing animal. Thereafter, we observed a decreased HR to a low-point at around P15, consistent with the widely recognized inverse relation of HR with age in human fetal, neonatal and infant periods^17^. This data contrasts with Wiesmann and colleagues who reported that HR measurements were similar over time, when measured in later stage neonates^16^.

The EF for the mice scanned at P1 (the earliest time point) was recorded at approximately 72% (Fig. 1d). This reduced to about 65% in 2-4 days, and then continued to reduce to the more adult heart-like value of approximately 60% during the following two weeks^14^. This data may reflect the many changes happening in heart development during the first week after birth, such as a reduction in relative cardiomyocyte proliferation and cell cycle exit^4^ and the rapid expansion of the coronary vasculature^18^.

### In vivo MRI

Next we analysed end-diastolic and end-systolic MRI cine images of P1 and P7 mice at baseline (Supplementary Fig. 2a, d), day 4 (Supplementary Fig. 2b, e) and day 21 (Supplementary Fig. 2c, f; Supplementary Movies 1-4), comparing MI and sham-operated groups in order to inform on the extent of injury and resolution over time. Day 21 was specifically selected as the time-point of reported complete regeneration following injury at P1^4^. Growth of the neonatal mouse heart was equivalent between MI and sham-operated animals for the P1 group (Supplementary Fig. 2a-c). We observed clear indications of tissue injury in the P1 mouse at day 4 post MI as indicated by regions of akinesis in the ventricle wall around the site of injury (Supplementary Fig. 2b), this was resolved by day 21 (Supplementary Fig. 2c; Supplementary Movies 1 and 2). Evidence of injury at the day 4 time point in P1 mice post-MI was confirmed by histology (Supplementary Fig. 3a-c). In contrast to P1 hearts, the P7 cohort following injury revealed significant dilation of the left-ventricle at days 4 (first scan after injury; Supplementary Fig. 2e) and 21 (experimental endpoint; Supplementary Fig. 2f and Supplementary Movies 3 and 4) compared to corresponding sham controls.

**Figure 3.**
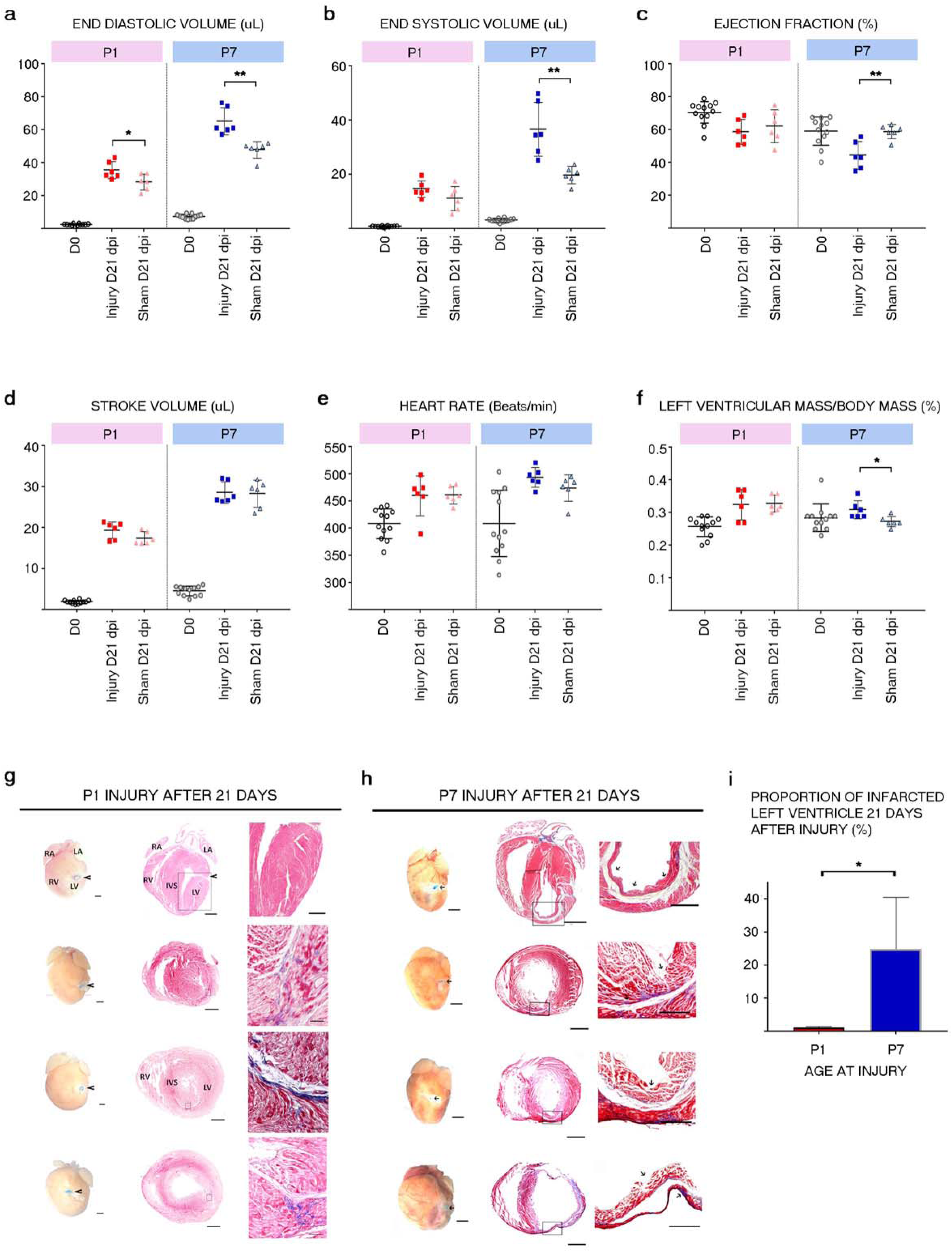
Left-ventricular cardiac functional parameters and representative histology at baseline (Day 0) and experimental endpoint (Day 21). End-diastolic volume (EDV) measurements revealed a more significant increase in EDV for the P7 injured cohort than the P1 cohort compared to matching sham controls (p < 0.0014 vs p < 0.034) suggesting increased ventricular remodeling to compensate for sustained myocardial damage. (b) End-systolic volume (ESV) revealed that only the P7 cohort showed a significant increase in ESV at 21 days (p < 0.0025). (c) Ejection fraction (EF): a significant decrease in EF for the P7 group (p < 0.0038) indicative of reduced heart function, whilst the injured P1 EF was comparable with that of matched sham controls. (d) Stroke volume (SV): both injury groups showed a similar SV to sham controls. (e) Heart rate (HR) measurements revealed an increase in HR for the P7 injury group relative to controls was observed but this was not significant (p = 0.14). (f) Left-ventricular mass index (LVM/BM) increased significantly (p < 0.019) for the P7 injury group relative to sham controls but there were no differences between P1 injured hearts and shams. The open circles indicate the baseline results (i.e. before intervention), the squares the injured hearts and the triangles the sham-operated animals at day 21 post intervention, respectively. Panels (g) and (h) show whole heart (left columns) and Masson’s trichrome staining of long axis (top middle and right) and short axis slices (remainder of middle and right columns) at different magnifications (boxed regions are higher magnifications, illustrated to the right), clearly illustrating the differences between injuries sustained at P1 and P7. The P1 hearts reveal mostly healthy and viable myocardium with minimal scarring (blue areas of collagen deposition) and thick ventricular walls. The P7 hearts show LV wall thinning and substantial collagen deposition in all hearts at all time points. The proportion of infarcted left ventricle after 21 days was quantified for both the P1 and P7 hearts (Panel I; * p <0.05). Scale bars: (g) left and middle columns = 500μm; right = 50μm. (h) left and middle columns = 500μm; top right = 100μm; rest of right column = 50μm.

**Figure 2.**
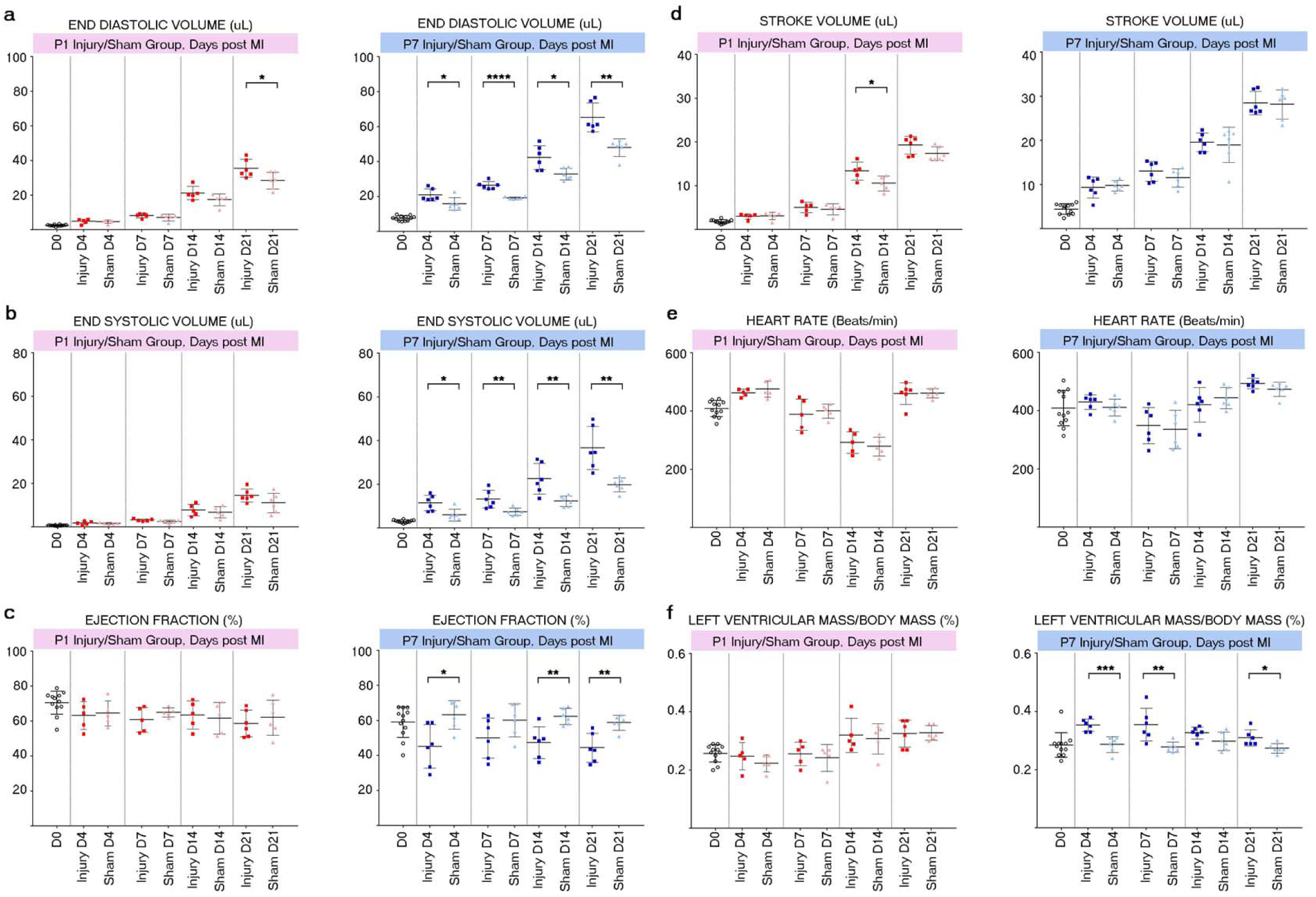
Functional MRI parameters at baseline and days 4, 7, 14 and 21 post-MI and sham-operated P1 versus P7 mice. MRI values for End Diastolic Volume (EDV; a), End Systolic Volume (ESV; b), Ejection Fraction (EF; c), Stroke Volume (SV; d), Heart Rate (HR; e) and Left Ventricular Mass to Body Mass ratio (LVM/BM; f) are indicated. Each time point shows a direct comparison between the injury and the sham groups within each P1 and P7 cohort; significant differences between injury and sham cohorts are indicated as *P < 0.05; **P < 0.01; ***P < 0.001 and ****P < 0.0001; n=5 or 6 for all groups.

These data collectively revealed significant differences in recovery between injured P1 and P7 hearts over 21 days post-MI, that manifested as pathological remodeling and abnormal left ventricular function in P7 hearts post-MI which was reversed and restored respectively in animals injured P1.

### Left-ventricular functional parameters at baseline (day 0), day 4, day 7, day 14 and day 21 for MI injury and sham-operated mice

We performed quantitative analyses of cine MRI images to measure left-ventricular function at baseline (before surgery – day 0) and days 4, 7, 14 and 21 post-MI for both P1 and P7 age-groups versus matched sham-operated controls. We assessed left-ventricular end-diastolic (EDV), end-systolic (ESV), ejection fraction (EF), stroke volume (SV), heart rate (HR) and left ventricular mass/body mass index (LVM/BM; Figure 2). EDV for the P7 injured group was significantly higher than that for the sham controls at all time points post-injury (Fig. 2a; Day 4: Injury 20.9 ± 1.4 vs Sham 15.8 ± 1.5, P < 0.05, n=6; Day 7: Injury 26.4 ± 0.9 vs Sham 19.1 ± 0.3, P < 0.0001, n=6; Day 14: Injury 42.3 ± 2.8 vs Sham 32.8 ± 1.4, P < 0.05, n=6; Day 21: Injury 65.2 ± 3.4 vs Sham 48.0 ± 2.1, P < 0.01, n=6;) indicative of ventricle dilation and remodeling. For the P1 group, there was no difference immediately after injury, but a rise in EDV in the injury group was observed, which became significant by day 21 (Injury 35.6 ± 2.1 vs Sham 28.5 ± 2.0, P < 0.05, n=6; Fig. 2a) inferring that regeneration may be incomplete by this time-point in contrast to previous reports^4, 8^.

ESV values for the P7 injured group were also significantly higher across all time points (Day 4: Injury 11.5 ± 1.5 vs Sham 6.0 ± 1.1, P < 0.05, n=6; Day 7: Injury 13.3 ± 1.7 vs Sham 7.6 ± 0.7, P < 0.01, n=6; Day 14: Injury 22.6 ± 2.9 vs Sham 12.4 ± 1.0, P < 0.01, n=6; Day 21: Injury 36.7 ± 4.0 vs Sham 19.8 ± 1.3, P < 0.01, n=6; Fig. 2b), such that by day 21 the ESV was almost twice that of sham controls. The P1 injured group revealed small relative increases in ESV, which were not significant (Injury 14.6 ± 1.2 vs Sham 11.1 ± 1.9, P = 0.15, n=6; Fig. 2b).

The ejection fraction (EF) was significantly lower for the P7 injury group, compared to controls, for most of the time points analyzed (Fig. 2c): there was a decrease in EF immediately post-MI, an apparent recovery at day 7 but subsequent further significant decreases in EF through days 14 and 21, suggesting impaired recovery of normal heart function over time (Day 4: Injury 45.3 ± 5.1 vs Sham 63.3 ± 3.3, P < 0.05, n=6; Day 7: Injury 50.1 ± 4.7 vs Sham 60.3 ± 3.9, P > 0.05 (not significant), n=6; Day 14: Injury 47.4 ± 3.6 vs Sham 62.4 ± 1.9, P < 0.01, n=6; Day 21: Injury 44.5 ± 3.4 vs Sham 58.8 ± 1.8, P < 0.01, n=6). In the P1 group there were no significant differences between the injured and sham hearts at all time points (Fig. 2c); the EF was maintained at around 60%, as previously reported for intact neonatal mice^16^.

Stroke volume (SV) increased linearly for both P1 and P7 hearts as the mice aged; the only significant difference between injury and sham being an increase in the P1 group at day 14 post-MI (Injury 13.4 ± 0.9 vs Sham 10.6 ± 0.8, P < 0.05, n=5; Fig. 2d).

There were no significant differences in heart rate (HR) between the injured P1 and P7 versus sham controls over the duration of imaging (Fig. 2e). However, the HR changes over this time period were not linear and represented a sinusoidal curve from birth to P28, with a low point for both groups at around P14-P15. This differs from a previous study which indicated no difference in HR with age from 3 days to 16 weeks after birth, or even a slight increase for a second follow-up group^19^. In our study measurements of HR were initiated from an earlier P1 time point and across a more detailed time course which should more accurately reflect HR dynamics over this time period; including accommodating changes in neural activity (the developing sympathetic and parasympathetic nervous system) that influence HR during the immediate neonatal period^20^.

The ratio of left ventricular mass over body mass (LVM/BM) was significantly increased in the P7 group at days 4, 7 and 21 after injury (Day 4: Injury 0.35 ± 0.01 vs Sham 0.29 ± 0.01, P < 0.001, n=6; Day 7: Injury 0.36 ± 0.02 vs Sham 0.28 ± 0.01, P < 0.01, n=6; Day 14: Injury 0.33 ± 0.01 vs Sham 0.30 ± 0.01, P > 0.05 (not significant), n=6; Day 21: Injury 0.31 ± 0.01 vs Sham 0.27 ± 0.01, P < 0.05, n=6; Fig. 2f), consistent with an adult cardiomyocyte hypertrophy response, whilst no significant differences were observed between the P1 injured versus sham controls (Fig. 2f). The previous intact MRI study revealed that LVM/BM decreased with increasing age at later stages^16^ but the situation appears more complex during the first three weeks of growth: there was a reduction in LVM/BM immediately after birth during the first week, then an increase for the second week, followed by a further decrease after three weeks. This suggests that around 14 days post-MI (P15) there is a compensatory effect to increase LVM. This is consistent with a report that between P10 and P16 at baseline (no-injury) heart growth exceeds that of the body, and subsequently between P16 and P21, body growth exceeds that of the heart; which in turn has been attributed to a T3-hormone induced proliferative burst of cardiomyocyte numbers during the early preadolescence phase at P14^21^.

These data collectively reveal pathological remodeling of the P7 heart post-MI which was significantly reduced and transient in the P1 heart following injury; consistent with restoration of cardiac output and the increased regenerative capacity of the latter. This was further supported by the different responses in the injured P1 versus P7 hearts at Day 21 post-MI (Figure 3). This endpoint data also revealed that, while the P1 injured neonate heart recovers the majority of function with similar values for all the measured cardiac parameters as matching sham-injured control hearts, there was some residual remodeling, as indicated by significantly higher EDV (Fig. 3a) that remained even after 21 days.

### Left-ventricular function over time following injury

In order to further elucidate the evolution of left ventricular function post-surgery in both age groups, we quantified EDV, ESV, EF and LVM/BM over time in the same mouse (Figure 4). In P1 mice these functional parameters were preserved in injured hearts relative to sham controls up to 21 days post-MI, with increases over the time course in both MI and sham groups reflecting heart growth (Figure 1). In contrast, we observed significant differences in functional parameters between injured and sham hearts for the P7 group from day 4 post-injury until day 21. Notably, differences in EDV (Fig. 4a) and ESV (Fig. 4b) between MI and sham were most pronounced for the P7 group, indicative of elevated pathological remodeling and which coincided with reduced function of the heart as reflected by a significantly lower EF at days 4, 14 and 21 post MI (Fig. 4c) also clearly visible in the MRI cine images of Supplementary Fig. 2b-c, e-f). These findings were consistent with our histological data (from a different cohort of mice) post-MI, whereby short-axis (Supplementary Fig. 3a-b – Day 4; Fig. 3c – Days 7, 14 and 21) and long-axis (Fig. 3g, h – Day 21) sections revealed a restored myocardium following transmural infarct in the left ventricle in P1 neonates versus persistent scarring and chamber dilatation in the P7, indicative of regeneration in the P1 but not the P7 group (Fig. 3g, h and Supp. Fig. 3a, b). The extent of initial scarring at Day 4 (Supplementary Fig. 3b) and persistent scarring by Day 21 (Fig. 3i) in the P7 injured heart versus P1 was significant in both cases (P < 0.05).

**Figure 4.**
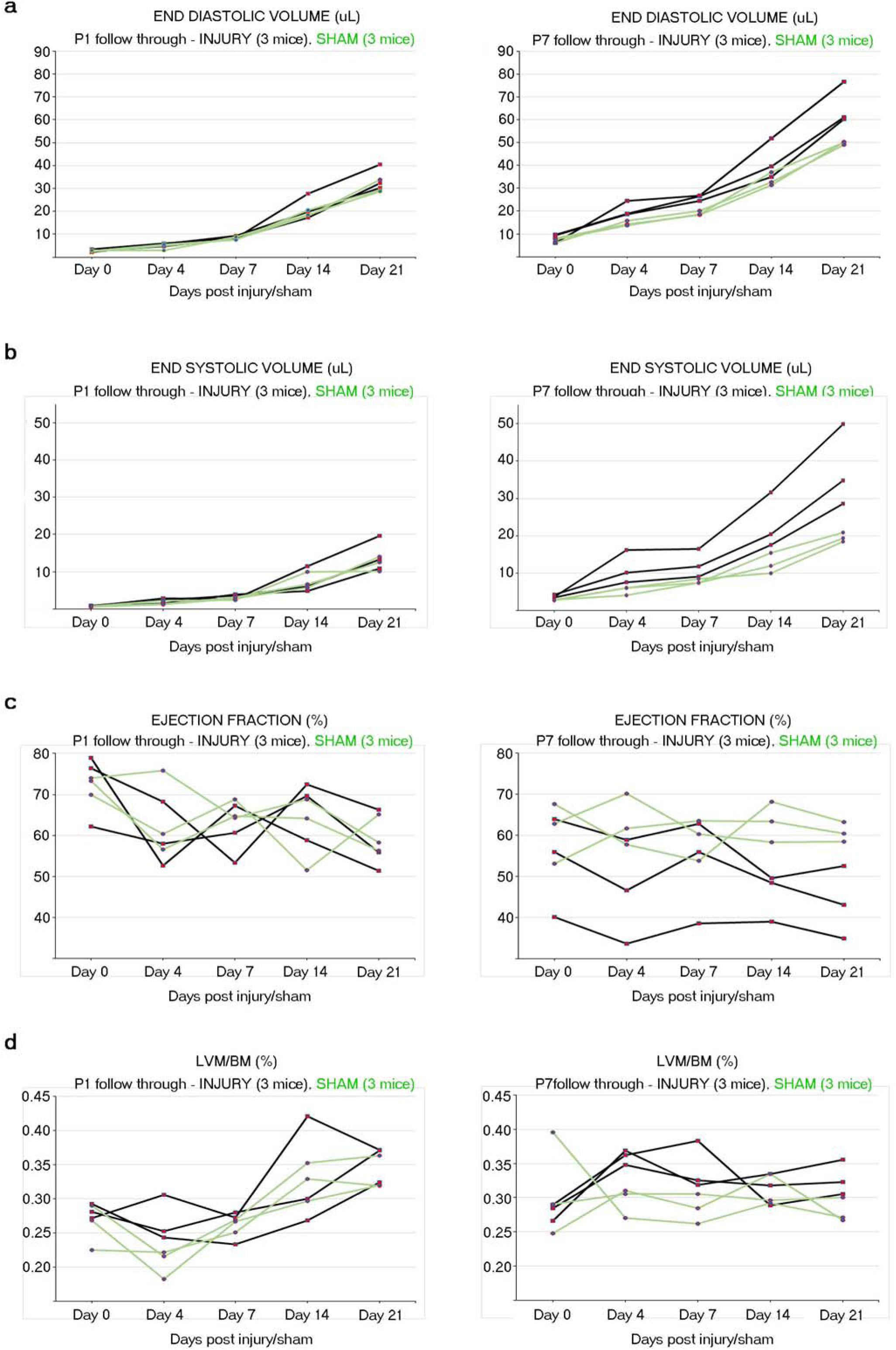
Longitudinal imaging of individual neonatal mice across five time points from baseline (Day 0) through to the study endpoint (Day 21). Development of (a) EDV, (b) ESV, (c) EF and (d) LVM/BM at Days 0 (baseline), 4, 7, 14 and 21 (post-MI) for P1 (left column) and P7 (right column) cohorts. Values obtained in sham-operated animals are shown in green and the ones from MI injured hearts in black, respectively (n=3 for each group). Clear differences in all these cardiac function parameters are evident for the P7 injured group from Day 4 through to Day 21, illustrating remodeling and reduced heart function. The P1 group more closely resembled their sham-operated control set, indicative of restored cardiac output, reversed remodeling, coincident with tissue regeneration.

In **summary**, we present a novel and significant adaptation of MRI to longitudinally assess tissue regeneration and cardiac function in a living mammalian model of heart regeneration. This provides definitive evidence for regeneration of the heart in the P1 mouse following injury and important insight into the output and remodeling of the regenerating heart over time, as directly compared to P7 animals which undergo a default adult wound-healing response. The regenerative capacity of the injured P1 mouse heart may simply be explained by harnessing the intrinsic growth responses that are ongoing following birth, however, the mechanisms of tissue growth and replacement following injury are highly likely to be relevant for targeting in the adult situation.

The neonatal mouse MRI method described can be applied by all groups with small animal MRI capability, and delivers a platform for future testing of therapeutic agents to enhance myocardial regeneration for ultimate extrapolation to the diseased adult human heart.

## Methods

### Animal husbandry and maintenance

Male and female CD1 mice were obtained from a commercial breeder (Harlan, UK) and kept under controlled conditions for temperature, humidity, and light, with chow and water available ad libitum. Typically, breeding trios (i.e. 1 male with 2 females) were setup to generate the pups for the P1 and P7 imaging studies. The female mice were plug-checked daily to estimate date of pup delivery. The mice were housed in a dedicated animal care facility with a controlled environment (12 hour light/dark cycle, 20°C, 50% humidity). Food (regular mouse chow) and water were available ad libitum, and the cage bedding was changed regularly until a few days before anticipated birth of a litter. At that time, the males were removed from the cage, and the pregnant females separated into individual cages to litter down. Pups were housed with the mother until weaning.

### MR hardware

All experiments were carried out on a 9.4 T (400 MHz) MRI system (Agilent Technologies, Santa Clara, CA) comprising a horizontal magnet (bore size 210 mm), a VNMRS DirectDrive^TM^ console, and a shielded gradient system (inner diameter (i.d.) 60 mm, gradient strength 1 T/m, rise time 130 µs). Purpose designed mouse cradles (with outer diameters of 20 mm for P1-P4 mice, 28 mm for P7-P14, and 33 mm for pups older than P14 – Fig. 3 A) including anaesthetic nose cone were either custom made from acrylonitrile butadiene styrene (Solidworks, Dassault Systèmes Solidworks Corp., USA; HP Designjet 3D-printer, Hewlett-Packard, USA) or sourced from the manufacturer (Agilent Technologies).

### Animal preparation for imaging

Pups were housed with the mother until weaning. Prior to examination by MRI, all pups were simultaneously removed from the mother and placed together in a heated enclosure at 32°C. In order to prepare for cine-MRI, anesthesia was induced in an anesthetic chamber using 4% isoflurane in 100% oxygen. Animals were weighed and then positioned prone in the cradle and maintained at 1.5%-2.0% isoflurane at 1.5 L/min oxygen flow throughout the MRI. A custom-made heated air blanket was calibrated to yield a surface temperature of 36°C, and was placed on the back of the pups to maintain their body temperature. A miniature pressure pad made from polyurethane foam and polyvinylchloride film fixed to a polythene tube was placed beneath the pups, and connected to a commercial pressure transducer (RS Components, Corby, UK) to derive respiratory and heart rate signals of the pups in the magnet. Signals were continuously monitored using an in-house developed gating device^22^. The experimental setup is shown in Figure 1 with representative physiological monitoring traces. Once the scan was completed, the pup was removed from the cradle and allowed to recover in the warming chamber with its littermates. Upon completion of all experiments on a litter, the pups were covered with a mash of faeces collected from their home cage (to disguise external odors and thereby facilitate acceptance) and collectively returned to the dam.

### Myocardial infarction surgery

After performing the initial baseline scan (at day 1 or day 7), the pups were removed from the cradle and allowed to recover for 10-15 mins depending on the imaging duration. Whilst still under the influence of residual isoflurane, the pups were placed on ice for 1 (P1) or 2 minutes (P7) to induce a surgical plane of anaesthesia by hypothermia. They were then taped onto a frozen block and the chest cavity was carefully punctured and opened between two ribs on the left side. The heart was extruded by applying gentle pressure below the diaphragm and on the right side of the chest. The left anterior descending artery was ligated by placing a suture through the left ventricle wall and securing it. The heart was repositioned within the chest, the ribs sewn together and the skin sutured. The pup was removed from the ice-block and warmed rapidly in front of a heat lamp, with care being taken not to overheat. To aid recovery, extra oxygen was administered, and physical stimulation was provided by holding an active littermate in close proximity. When the pup was fully recovered, it was placed back in the recovery chamber with the rest of the litter. The individual steps of the surgical procedure are illustrated in Supplementary Fig. 4.

All investigations conformed to Home Office Guidance on the Operation of the Animals (Scientific Procedures) Act, 1986 (HMSO) and to institutional guidelines.

### MRI acquisition

After scouting for the short-axis orientation of the heart using a fast, spoiled 2D gradient echo sequence (TE/TR = 1.15 / 5 ms, flip angle 15°, number of averages 8, slice thickness 0.5 / 0.7 mm, field-of-view (FOV) 15 × 15 mm^2^ to 25 × 25 mm2), B0-maps were acquired (TE/ΔTE/TR = 0.86 / 0.75 / 5 ms, flip angle 15°, number of averages 4, slab thickness 11.2 to 15.0 mm, field-of-view (FOV) 20 × 20 × 15 mm^3^ to 25 × 25 × 20 mm^3^) to allow for automated shimming of a 3D volume across the cardiac region of mouse. Retrospectively gated, compressed sensing (CS) accelerated multi-frame imaging (‘cine-MRI’) was performed covering the heart from base to apex in a contiguous stack of short-axis slices (TE/TR = 1.86 / 4.6 ms, flip angle 10°, number of averages 1, number of frames 50, number of repetitions 6, slice thickness 0.5 / 0.7 mm, number of slices 8 - 12, field-of-view (FOV) 15 × 15 mm^2^/ 20 × 20 mm^2^). Each repetition was two-fold under sampled in phase-encoding direction and corresponding sampling patterns were generated as reported previously^23^. All imaging was performed without prospective cardiac and respiratory gating. The total imaging time (including experimental preparation) was 35-45 mins, primarily depending on the number of slices required to fully cover the left ventricle. Imaging was performed at before and days 4 and 21 post-surgical intervention.

### MRI data reconstruction

Reconstruction of the data was performed offline using a custom-written software tool ‘*bmru_itools’* developed in-house in *Interactive Data Language (*IDL 8.3; Harris Geospatial, Wokingham, UK). Analysis of the navigator echo was performed as described previously^24^ to yield *k*-space signals for 20 frames per slice, which were then subjected to CS-reconstruction as reported previously^25^, followed by isotropic zerofilling (factor of two), filtering (modified third-order Butterworth filter), and Fourier transformation. Examples for respiratory and cardiac traces obtained from the navigator signal are shown in Fig. 1b.

### Calculation of left-ventricular function

Epi- and endocardial borders were manually outlined in the end-diastolic and end-systolic frames (see Fig. 1d) to assess left-ventricular structural parameters (i.e. left-ventricular mass, LVM; end-diastolic volume, EDV; end-systolic volume, ESV), using the same *IDL* software tool. This allowed for the calculation of the left-ventricular functional parameters stroke volume (SV = EDV – ESV) and ejection fraction (EF = SV / EDV · 100%). LVM is reported as the mean of the respective end-diastolic and end-systolic volume multiplied by the specific weight of cardiac muscle tissue of 1.05 g /cm^3^.

### Statistical analysis

Animal numbers and sample sizes reflected the minimal number needed for statistical significance based on previous experience. MRI data analysis was performed offline after all scans were complete to blind researchers to initial surgical procedure. Statistics were calculated using Microsoft Excel and Prism Graphpad software. The statistical significance between two groups was determined using unpaired two-tailed Student’s *t*-test and the data were reported as mean ± s.e.m, with n=6 for all groups. A value of *P*<0.05 was considered statistical significant.

### Tissue collection

Mice were euthanized by hindbrain destruction, hearts were excised and washed in ice-cold PBS prior to fixation in 4% paraformaldehyde overnight at 4°C.

### Tissue preparation for staining

Hearts were dehydrated through an ethanol series (50%; 70%; 80%; 90%; 96%; 100%; 100%; at least 2 hours per solution) and were placed in butanol overnight. Samples were then washed in 50:50 butanol and paraffin wax (Histoplast, Fisher Scientific) at 56°C for one hour after which they were washed through 100% paraffin (56°C) twice (1 hour each time) before tissue orientation and embedding (in Histoplast). Sections were cut at 8μm on a microtome and mounted on Superfrost slides. Slides were left to dry overnight on a heated rack (40°C) before use. Sections were deparaffinised by washing through Histoclear (Fisher Scientific, twice for 5 minutes) and were rehydrated through ethanol series (as for dehydration but in reverse order).

### Haematoxylin and Eosin (H&E) staining

This protocol was performed at room temperature. Post rehydration, slides were washed once in PBS before being immersed in Haematoxylin (Sigma Aldrich, UK) for 5 minutes. The slides were then cleared under running tap water for 5 minutes (or until the water ran clear). Following immersion in distilled water, slides were immersed in Eosin staining solution (Sigma, UK) for approximately 1 minute and then rinsed in distilled water. The slides were then quickly dehydrated (as for rehydration, in reverse order) and mounted using DPX mounting media (Sigma, UK).

### Masson’s trichrome staining

This protocol was performed at room temperature according to manufacturer’s instructions (Sigma Aldrich, UK; HT-15). Briefly, post-rehydration, slides were washed once in PBS before being immersed in Bouin’s fixative solution overnight. Slides were then immersed in the following solutions: Haematoxylin (5 minutes then wash under running tap water followed by distilled water); Beibrich Scarlet-Acid Fushchin (5 minutes then wash with distilled water); Working-Phosphotungstic/Phosphomolybdic acid solution (5 minutes); Analine Blue solution (5 minutes); 1% Acetic Acid solution (30 seconds). Slides were then rinsed in distilled water and quickly dehydrated (through ethanol series into Histoclear) and were mounted using DPX mounting media.

## Acknowledgements

This work was generously supported by the British Heart Foundation via a chair award (CH/11/1/28798; PRR), and senior fellowship (FS/11/50/29038; JES) and through the BHF Oxbridge Regenerative Medicine Centre (MGR, JES and PRR; RM/13/3/30159); and an EU Marie Curie FP7 Innovative Training Network: CardioNet. (MM; #289600). The authors would like to thank the late Ms Victoria Thornton for supporting the MRI work, and Dr Tobias Wech (University of Würzburg, Germany) for providing the optimized Compressed Sensing sampling schemes. We also appreciate the technical and administrative support from Ms Lee-Anne Stork and Ms Aude Vernet during the surgery and scanning procedures.

## Author contributions

PRR and JES conceived and designed the original study. MM carried out pilot MRI studies and the histology. MLM, SCS and JES designed and built the custom hardware for neonate MRI scanning. JES developed the data acquisition method and the image analysis software. MGR performed the neonate surgeries and MRI scanning with MLM. MGR analyzed the data and prepared it for publication. The manuscript was written by PRR, MGR and JES. All authors approved the final version of the manuscript.

## Additional information

### Competing interests

PRR is co-founder and equity holder in OxStem Cardio, an Oxford University spin-out that seeks to exploit therapeutic strategies stimulating endogenous repair in cardiovascular regenerative medicine.

